# Some rats take more time to make easier perceptual decisions

**DOI:** 10.1101/2021.10.26.465587

**Authors:** Pamela Reinagel

**Affiliations:** University of California San Diego, La Jolla CA 92093 USA

## Abstract

When subjects control the duration of sampling a sensory stimulus before making a decision, they generally take more time to make more difficult sensory discriminations. This has been found to be true of many rats performing visual tasks. But two rats performing visual motion discrimination were found to have inverted chronometric response functions: their average response time paradoxically increased with stimulus strength. We hypothesize that corrective decision reversals may underlie this unexpected observation.

## Background

In some studies of two-alternative forced-choice (2AFC) sensory discrimination, subjects are given unlimited time to sample the stimulus before responding, with no time limit or response cue. In such experiments in rodents, response times have been reported to be longer for more ambiguous visual discriminations, for example in auditory click rate discrimination (*1*), random dot motion discrimination (*2*), morphed image discrimination (*3*), olfactory discrimination (*4*), visual orientation discrimination (*5-7*) or looming visual stimulus escape responses (*8*). This is as expected from the previous literature on human and monkey decision-making, and as predicted by a drift diffusion model (DDM) for perceptual decisions (*9*). Recently, however, we observed two rats with markedly inverted chronometric response functions: response time increased with increasing stimulus strength. Here we report this observation and suggest an explanation.

## Methods

No new experiments are reported in this paper; the data are from a previous publication (*10*) and the previously unpublished supplemental video is from (*2*). Those studies were both performed in full accordance with ethical guidelines for animal research, as approved and overseen by the UCSD IACUC. Rats earned all their daily water through the task. To briefly summarize the behavioral task: the stimulus was a field of randomly scattered white dots displayed on a black background. The “signal” dots moved at a fixed velocity either to the left or right side. The “noise” dots moved at random velocities. The strength of the motion signal was controlled by the “coherence”, defined as the fraction of dots that were signal dots. The size, contrast, number, and speed of the dots were selected to ensure that accuracy ranged from 50% (chance) to 100% correct for most rats. Rats freely initiated each trial; stimuli appeared without delay and continued until the rat responded either Left or Right. Responses that correctly identified the direction of the coherent motion were rewarded with water. The direction and coherence were randomly selected independently each trial. In the no-signal (0% coherence) condition, the “correct” response was randomly assigned. Of particular relevance to this analysis, the task imposed no minimum response time, no minimum time to reward, no post-response reward delay, and no enforced inter-trial interval. Based on the observation that most responses were made in <3s, the stimulus duration was experimentally limited to either 3s or 5s, and for this reason trials with RT>3s were excluded from analysis. Otherwise, here we include all available trials after task acquisition, which we defined as the earliest time point with >80% correct responses on an easy coherence for >200 consecutive trials.

In Fig. 1 E-H a two-tailed Spearman’s correlation was used because both underlying distributions are non-Gaussian. Any *p* values less than 1e-4 are reported as inequalities because lower values are not reliable. The comparisons made in Figs. 2 and 3 were not premeditated, but rather explorations in response to the unexpected observation in Fig. 1. Therefore, no claim is made or implied about statistical significance of these trends and no *p* values are reported. All data and code required to replicate the figures in this report are available in a replication-certified repository (*11*).

**Figure 1.**
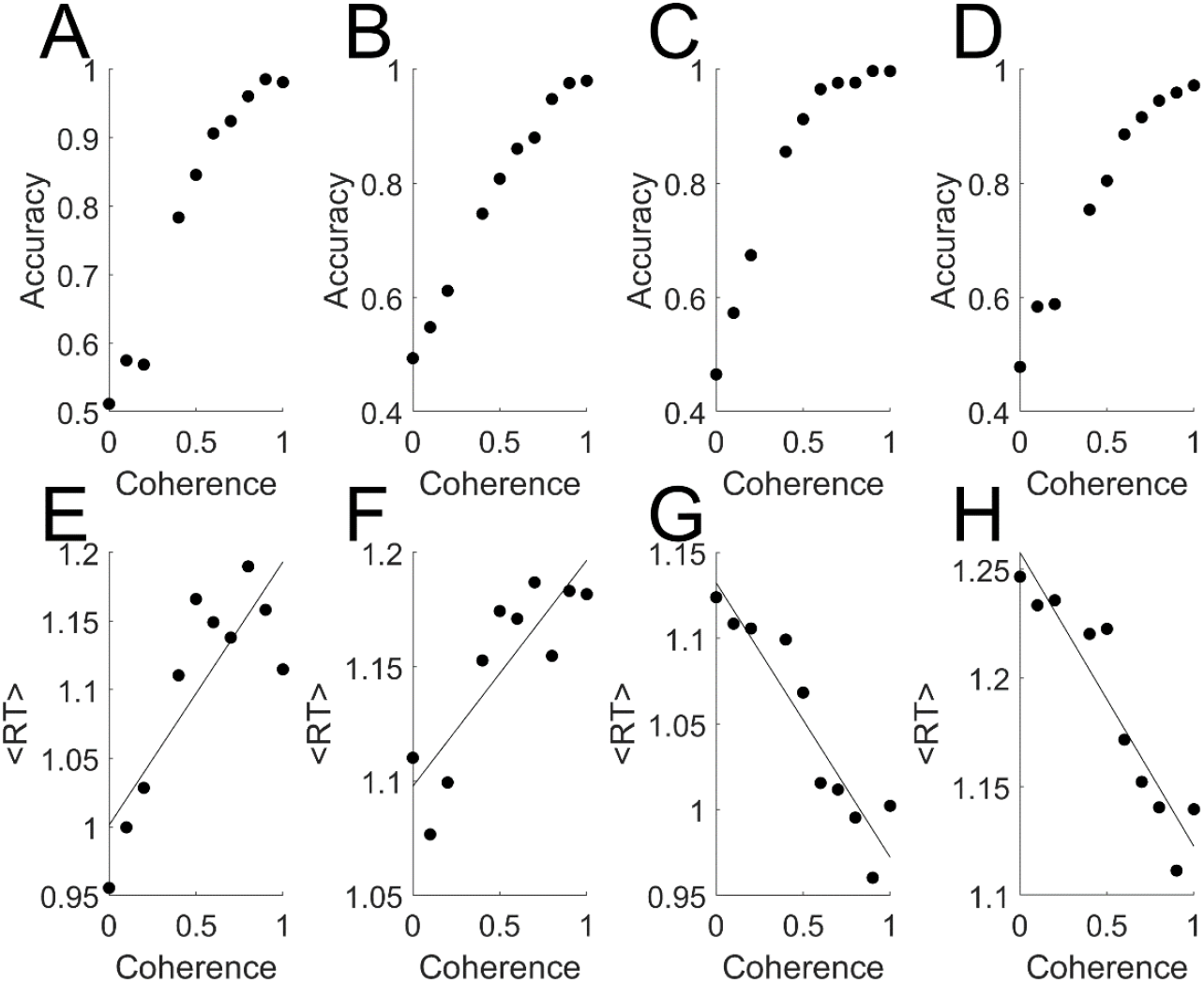
Rats with inverted chronometric response functions. Data from four rats in the same cohort, all performing a random dot motion task with randomly interleaved coherence values. **A-D**: Psychometric curves ranged from chance (50%) to near 100% in each case. **E-H**. Chronometric response functions from the same rats and trials analyzed in A-D. **E**. N=5396 trials; r = +0.71, *p* = 2.75e-2. **F**. N=6602, r = +0.81, *p* = 8.24e-3. **G**. N=4897, r = −0.96, *p* <1e-4. **H**. N=7223, r = −0.96, *p* <1e-4.

**Figure 2.**
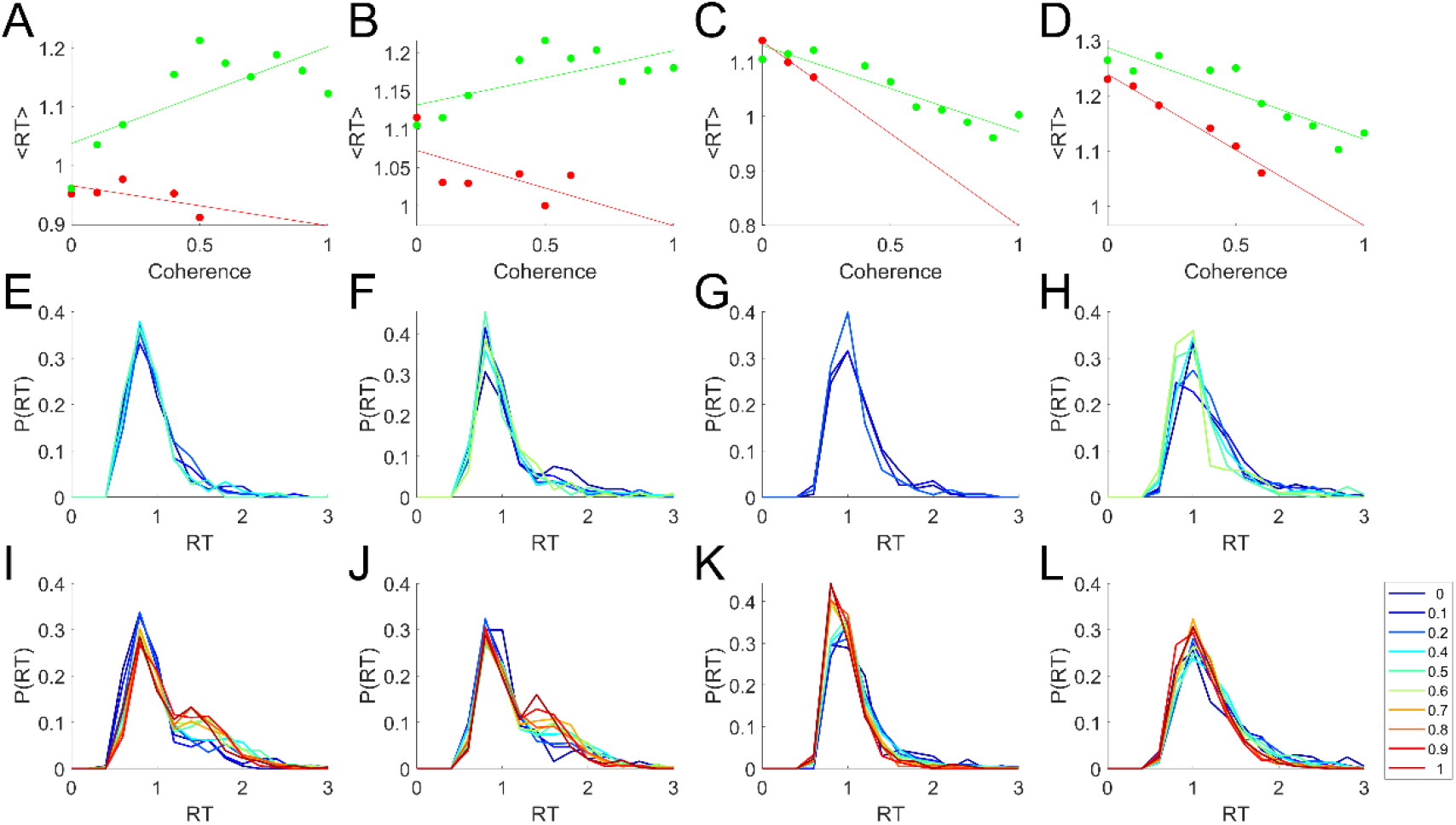
Inverted chronometric response functions are associated with bimodal correct trial RT distributions. Analysis of same data as Figure 1. A minimum of N=100 trials were required to estimate the mean or response time distribution for any coherence and trial outcome. **A-D**. Mean response time vs. stimulus coherence computed separately for correct trials (green) and error trials (red). Least square regression lines are also shown. **E-F**. Response time distributions of the error trials from A-D expressed as probability distributions (each curve sums to 1); color indicates stimulus coherence (legend at bottom right). **I-L**. Response time distributions of the correct trials from A-D.

**Figure 3.**
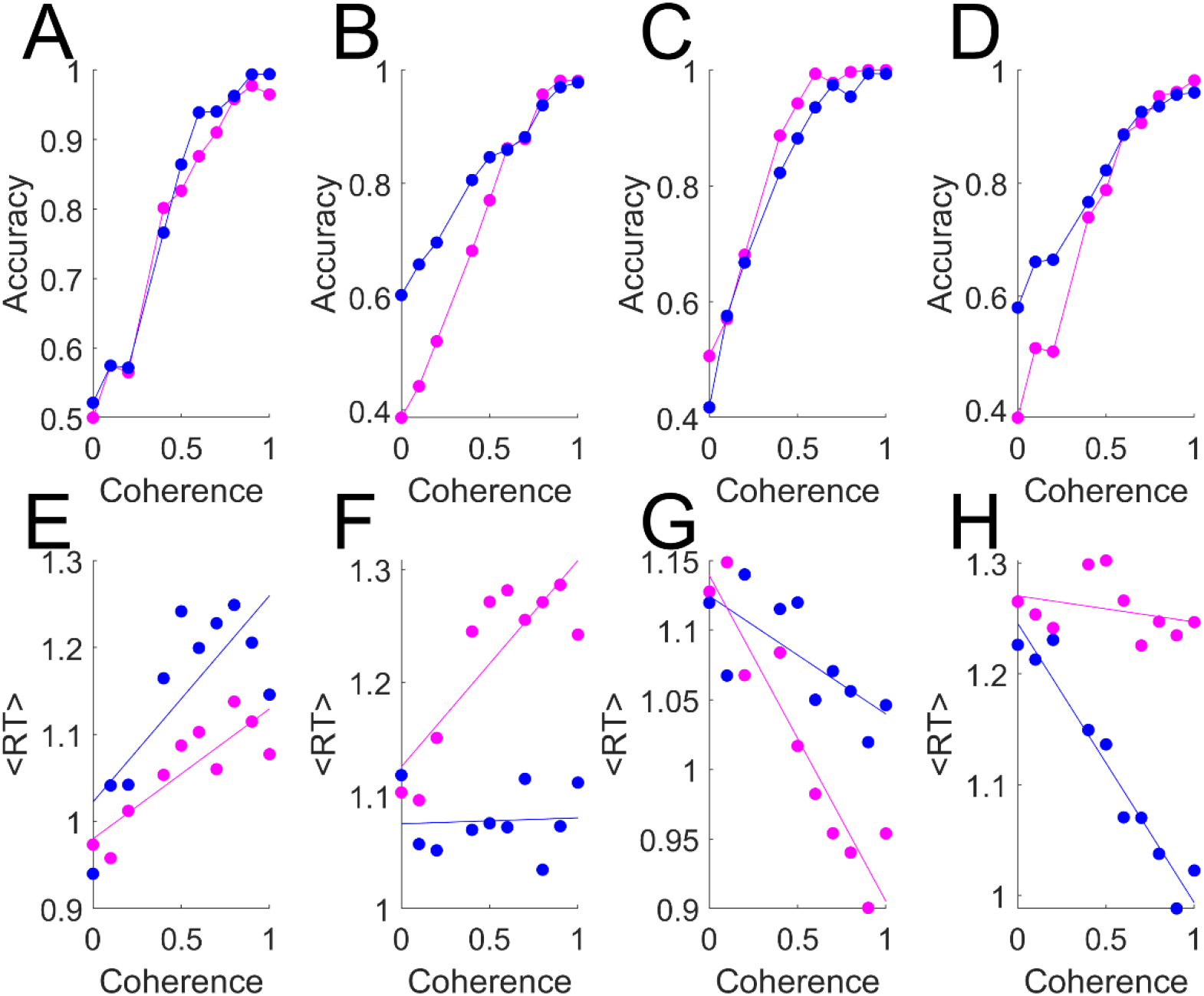
Psychometric and chronometric response functions according to response target. Analysis of same data as Figures 1-2. Trials whose target was the Left response port (magenta) vs. Right port (blue) analyzed separately. **A-D**. Accuracy of responses as a function of motion coherence (stimulus strength). **E-F**. Mean response time as a functin of motion coherence, with least square regression lines.

## Results

We noticed with surprise that some rats in a previous study (*10*) had longer mean response times to easier random dot motion stimuli (Figure 1 E,F), in spite of having unremarkable psychometric curves with minimal lapse (Figure 1A,B, performance at high coherence near 100%). Other rats in the same cohort with comparable psychometric curves (Figure 1 C,D) had the expected, decreasing chronometric response function (Figure 1 G,H).

The inverted chronometric response functions were attributable mostly to the correct trials (Figure 2 A,B green; c.f. A,B red or C,D green). The response time distributions of these rats revealed a prominent shoulder or second peak in the response time distribution in the correct trials (Figure 2I-J) that was absent or less prominent among error trials (Figure 2E,F) or rats with typical chronometric response functions (Figure 2 K,L). To the extent this could be resolved, the late response peak in the correct response time distributions increased with coherence (Figure 2I,J: above ∼1.2 s, warm colors are above cool colors), whereas in error trials (Figure 2E-H) or correct trials of typical rats (Figure 2K,L), we see heavier tails for lower coherences as expected from a diffusion to bound decision process (above ∼1.2 s, cool colors are above warm colors). We hypothesize that the late peak in the response time distributions of the unusual rats (Figure 2I,J) represent corrective decision reversals.

One of the rats with an inverted chronometric response function had only a slight response bias (Figure 3A); its chronometric response function was inverted for both the Left-target and Right-target trials (Figure 3E), consistent with the interpretation that decision reversals occurred for both targets. The other rat had a marked response bias at low coherence, which was overcome at high coherence (Figure 3B). In this case the chronometric response function was inverted mostly for the trials with the non-preferred target, and non-preferred side responses were also slower than preferred side responses. This is consistent with the idea that the late trials occurred primarily when the rat began to respond toward the preferred side, but reversed this plan later on the basis of stimulus evidence for the non-preferred side.

The example rats with ordinary chronometric response functions also had response biases (Figure 3C-D). These rats had decreasing response time for increasing stimulus strength for both Left and Right targets (Figure 3G-H), although the response times were longer and the slopes of the chronometric response functions were shallower for the non-preferred targets.

## Discussion

Early in a trial, animals could begin covert motor planning, adopt an oriented posture, or start moving toward one response on the basis of either non-stimulus information or early stimulus sampling. If the stimulus continues to be sampled, however, subsequent stimulus information could override the initial choice and the rat could abort or reverse the motor plan, resulting in a correct but delayed response. Stronger visual stimuli are more likely to accumulate sufficient evidence to the contrary in time to override an incorrect initial motor plan, and therefore would produce more corrective decision reversals. If such an effect were sufficiently pronounced, inversion of the chronometric response function could occur. The rats we observed with inverted chronometric response functions achieved near-perfect performance on high-signal stimuli, but used extra time to do so.

In the experiments analyzed here, we cannot directly test the decision-reversal hypothesis because there are no data on the rat’s behavior prior to decision commitment (but see Supplemental Video for an anecdotal example trial). Other tasks that use continuous report modalities (such as a joystick, trackball, wheel turning, or video analysis; or in the case of primates, gaze tracking) can provide direct evidence of decision reversals (*1, 5*).

As an aside, we note that if late-response trials are enriched for reversals from errors to correct decisions within any given coherence, this would provide an additional or alternative mechanism for the increase in accuracy with response time noted previously (*2, 3, 10*). Nevertheless, the inverted chronometric response functions reported here are the exception. Many rats exhibit expected chronometric response functions and a single-peaked response time distribution, and these rats still show an increase in accuracy with response time (*2, 10*). Therefore, that distinct phenomenon likely involves other mechanisms as well (*12*).

Not many rodent perception studies use 2AFC tasks in which error and correct response time distributions are separately defined. Only a subset of those use tasks in which the stimulus strength varies and the sampling duration is controlled by subject response time. Even those studies don’t often report chronometric response functions. Therefore, we don’t know if inverted chronometric response functions are rare or common in rodent data. The effect came to our attention because these particular rats had very large fraction of substantially delayed responses, yielding a markedly inverted chronometric curve. The same effect could easily go unnoticed if the late events had been less prevalent or less delayed, such that the chronometric response function was only flattened rather than inverted, and the response time distributions merely widened without a resolvable second peak. Indeed, we cannot rule out that this is occurring in the “normal” rats shown here, as we do see a difference in average response times and chronometric slopes for preferred vs. nonpreferred targets (Fig. 3G,H), consistent with corrective decision reversals. Therefore, it would be of interest to investigate more fully the role decision reversals might play in determining response times in perceptual decision-making in rodents as well as in primates and other species.

## Acknowledgements

Carly Shevinsky provided technical support for animal training and care, and first noticed the inverted chronometric response functions of these subjects.

## Supplemental Video

http://www.ratrix.org/images/RatDotMotion.mp4

## Notes

### Competing Interest Statement

The authors have declared no competing interest.

